# Evolution of Hierarchy in Bacterial Metabolic Networks

**DOI:** 10.1101/118299

**Authors:** Aaron Goodman, Marcus Feldman

**Author notes:** ^*^Correspondence.

## Abstract

**Background:** In self-organized systems, the concept of flow hierarchy is a useful way to characterize the movement of information throughout a network. Hierarchical network organizations are shown to arise when there is a cost of maintaining links in the network. A similar constraint exists in metabolic networks, where costs come from reduced efficiency of nonspecific enzymes or from producing unnecessary enzymes. Previous analyses of bacterial metabolic networks have been used to predict the minimal nutrients that a bacterium needs to grow, its mutualistic relationships with other bacteria, and its major ecological niche. Using flow hierarchy, we can also infer the tradeoffs between growth rate and metabolic efficiency that bacteria make given their environmental constraints.

**Results:** Using a comparative approach on 2,935 bacterial metabolic networks, we show that flow hierarchy in bacterial metabolic networks tracks a fundamental tradeoff between growth rate and biomass production, and reflects a bacterium’s realized ecological strategy. Additionally, by inferring the ancestral metabolic networks, we find that hierarchy decreases with distance from the root of the tree, suggesting the important pressure of increased growth rate relative to efficiency in the face of competition.

**Conclusions:** Just as hierarchical character is an important structural property in efficiently engineered systems, it also evolves in self-organized bacterial metabolic networks, reflects the life-history strategies of those bacteria, and plays an important role in network organization and efficiency.

## Background

In characterizing bacteria, we seek to understand both their internal processes and how they interact with other species and their environments. Techniques in cell and molecular biology have been very helpful in revealing the inner workings of bacteria, but do not address the ecological context in which bacteria develop and live. Increasingly, metagenomic techniques are being used to simultaneously sequence all of the bacteria present in a given environment. However, these techniques can only provide limited information about particular species, where they are found, their relative abundances, and co-occurrence patterns.

By studying the structure and evolution of a bacterium’s metabolic network, we can move beyond correlational profiles to understand both the underlying pressures that have driven its evolution as well as the ecological role it occupies. A bacterium’s ability to reproduce depends on the efficiency of its metabolism, which we can study as a network of metabolites linked together by the enzymes that transform one metabolite into another [1]. The structure of these networks varies across the bacterial kingdom and reflects the environmental pressures that guide bacterial evolution. Thus, a bacterium’s metabolic network can be used to predict the minimal nutrients that it needs to grow, its mutualistic relationships with other bacteria, and its major ecological niche [2][3][4]. In describing metabolic networks, two types of hierarchies can be helpful: flow hierarchy and containment hierarchy.

Previous study of the hierarchical nature of metabolic networks has tended to focus on containment hierarchy, which represents the nodes in a network as being contained within modules, which themselves are contained within other modules, and so on, in a recursive fashion. For example, a containment hierarchy may be used to represent the organization of a firm, with divisions, departments, teams and individual employees. Applied to metabolic networks, such modules correspond to known pathways [11][12], and modular hierarchy has been hypothesized to increase evolvability of metabolism [13]. Simulations of Boolean logic networks have suggested that modularity evolves in changing environments, and it has been hypothesized that this would be reflected in bacterial metabolic networks[14][15]. However this has not been borne out [16]; differences in modularity of metabolic networks have been found to be moderately correlated with the phylogenetic divergence of the organisms, and there is a general trend of loss of modularity over evolutionary time due to the addition of peripheral pathways during niche specialization [17].

In this work, we focus on the heretofore neglected type of hierarchy: flow hierarchy. While containment hierarchy represents the organization of a network as a series of modules, flow hierarchy characterizes the way information moves throughout the network. Information is acquired at the lowest level of the hierarchy and transmitted to higher levels, where it is aggregated and passed upward; at the same time orders come from the top of the hierarchical network and are passed down to lower levels.

Flow hierarchy is used to describe networks in many fields, particularly in the study of information accrual networks. In engineering, information accrual networks are used in the design of control systems, and in the social sciences, they are used to study the organization of firms [5][6]. To use the example of the firm, flow hierarchy could be used to represent the movement of orders and responsibilities throughout the firm, as low-level employees report upwards to supervisors who aggregate reports for department heads and so on, while orders flow downward from decision makers to executors. As bacteria synthesize the complex molecules needed for survival, they reduce the overall entropy within the cell. Given the thermodynamic equivalence of entropy reduction and information accrual, this reduction can also be viewed as an increase of information. Thus we can use the metabolic network graph to study the flow of information through the cell.

Although flow hierarchy (hereafter referred to as hierarchy) has not been well studied in metabolic networks, it has been identified in a variety of other self-organized networks, including food webs, neural networks, and the transcription factor network in *D. melanogaster*, where the degrees of hierarchy were significantly higher than would be expected in a random network with the same degree distribution [7]. In studying hierarchy in metabolic networks, we are able not only to learn that hierarchy appears higher than would be expected by a random configuration, but also to assess adaptive benefits that hierarchy provides.

These findings show that hierarchy is common among self-organized networks, but they do not explain when hierarchical network organization would provide a selective advantage. A number of comparative approaches have been used to make inferences about the forces guiding the development of networks in other disciplines, and from them we can deduce some of the adaptive benefits that hierarchy provides. Simulated evolution experiments of Boolean logic networks have shown that the cost of maintaining links between nodes is the driving force in the emergence of hierarchy [19]. Hierarchical characteristics have also been shown to predict the costs of maintaining information-sharing relationships in emergent social networks and reflect the degree of market variability that supply chains may be able to withstand [21]. We know that bacterial metabolic networks face similar constraints. Maintaining catalytic abilities between metabolites incurs a cost either as a trade-off between specificity and efficiency, or from production and replication of unused enzymes [20][22]. We show that the strength of these constraints is correlated with degree of hierarchy in metabolic networks.

To measure flow hierarchy quantitatively, researchers commonly use the global reaching centrality (GRC), defined as the average difference between the maximum local reaching centrality (i.e. fraction of nodes in the network accessible by each node of the network) and the local reaching centrality [23]. In essence, GRC is a measure of heterogeneity in the flow of information throughout a network. For example, a dictatorial firm where the boss exerts great influence over the entire company while individual employees have little sway would have a higher GRC than a consulting firm run by a group of partners, each of whom oversees a small group of highly collaborative employees. Other, less widely used measures of flow hierarchy are an eigenvector centrality based method, the fraction of edges participating in cycles, or by decomposition into treeness, feedfowardness, and orderability [7][8][9].

Studying the evolution of containment hierarchy can tell us about the environmental contingencies that inform the evolution of metabolic networks (e.g. resource availability, temperature, environmental variation, etc). However, by studying flow hierarchy we can also infer the different growth strategies a bacterium may pursue, furthering our understanding of how it fills its ecological niche [13]. We employ the reverse ecology principle to understand how the hierarchical character of a metabolic network reflects the life-history strategy of a bacterium in relationship to the growth-yield tradeoff, as well as its environmental niche. As bacteria first adapt to new habitats they may develop novel metabolic functions, leading to an increase in hierarchy, since the new metabolic functions are added to the periphery of the network.

As those ecosystems evolve, however, some bacteria may adopt higher-growth rate strategies in response to increased competition, at the expense of efficiency and consequently metabolic network hierarchy. Since only some bacteria adopt such high-growth rate strategies, while others maintain higher degrees of hierarchy and efficiency, variance in hierarchy increases overall. Though both hierarchy and modularity correlate with bacterial specialization, we find that, contrary to the Boolean network simulations, there is little evolutionary relationship between the two, and that there is more conservation of hierarchy than modularity over time.

## Results and Discussion

### Networks

Networks were reconstructed from 2,935 bacteria species in the KEGG database. These networks were robust to misannotation of enzymes. In random perturbations of the metabolic network for *E. coli* with 10% of the reactions removed, 95% the networks had hierarchy scores within 12% of the true network, and with 10% of reactions reversed, within 6% of the true network.

Network sizes ranged from 76 to 1496 metabolites, with a mean of 848. The smallest was the obligate insect parasite *Nasuia deltocephalinicola* and the largest was the soil bacterium *Burkholderia lata.*

### Hierarchy

Hierarchy scores for the metabolic networks were calculated using the GRC hierarchy score [23]. The mean degree of hierarchy was 0.279, and ranged from 0.065, for the insect symbiote Candidatus *Nasuia deltocephalinicola*, to 0.385 for a *Blattabacterium* endosymbiont of *Nauphoeta cinerea*, an insect endosymbiote. The hierarchy score for *E. coli* strains was 0.269 (Figure 1). For comparison with a random network and real world networks, GRC hierarchy scores for an Erdős-Rényi random graph is 0.058, a scale-free network 0.127, and a tree 0.997, an estuary food web 0.814, and the neuronal network of *C. elegans* [23].

**Figure 1.**
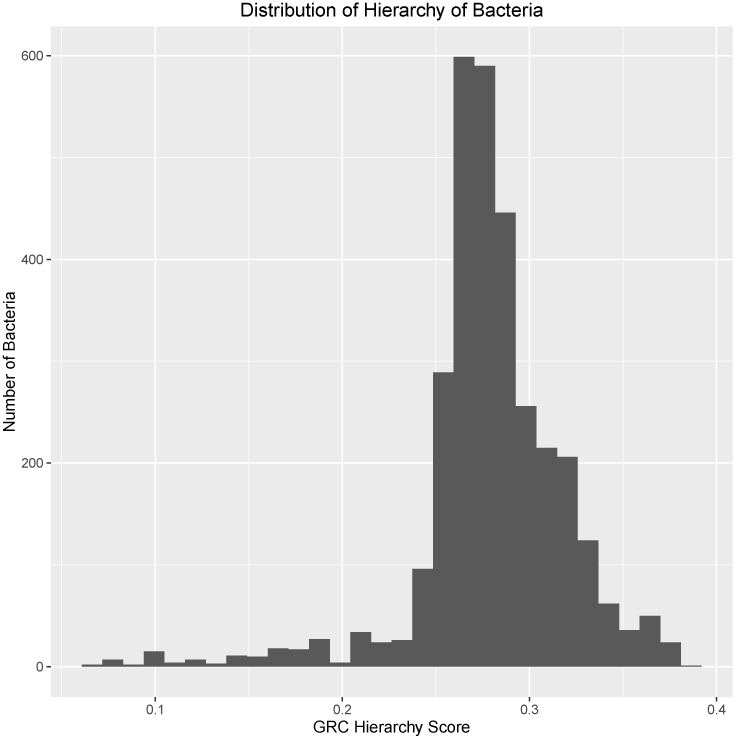
Histogram of GRC hierarchy scores of the 2,935 bacteria in the KEGG database. Mean degree of hierarchy is 0.279, ranging from 0.065 for the insect symbiote Candidatus *Nasuia deltocephalinicola* to 0.385 for a *Blattabacterium* endosymbiont of *Nauphoeta cinerea*, an insect endosymbiont. The hierarchy score for *E. coli* strains is 0.269.

### Relationship to environment and growth rate

There is a fundamental ecological trade off between growth rate and yield, which is a result of the underlying efficiencies of the reactions. Bacteria that have a metabolism that produces the maximal growth rate per amount of carbon taken up will have suboptimal biomass production, and vice versa.

This tradeoff is representative of fundamentally divergent ecological strategies that bacteria use [24]. Furthermore, the tradeoffs between growth and yield are represented in the constraints on the metabolic network, such that high-yield strategies lead to more hierarchical networks. There is a tradeoff between enzyme specificity and efficiency, so when yield is favored there will be higher costs of maintaining edges in the network, which leads to hierarchy [25] [20]. Rapidly growing bacteria have more metabolic cycles which allow for metabolic flexibility at the cost of wasted energy, and these cycles decrease hierarchy [26]. The cost of maintaining unused enzymes in the genome is higher when efficiency is paramount [22].

Using a dataset of 111 bacteria with known growth rates, we see that the hierarchical character of the network correlates inversely with growth rate, Spearman *ρ* = −0.31, *p* < 0.0007, fig 2. Furthermore, there is evidence that carbon efficiency constraints on bacteria differs greatly by environment, and that the evolutionary dynamics of carbon usage niche specialization are stronger within populations [27][28]. When we control for the bacterial environment, we see a correlation of *ρ* = −0.41, which is significantly greater than 0 (*p* < 0.0001, and significantly greater than the correlation when not controlling for the environment *p* < 0.003). Bacteria with hierarchy score greater than the median hierarchy score grow at a rate of 0.64 doublings per hour, compared to 1.44 doublings per hour for bacteria with hierarchy score less than the median, *i.e.* bacteria with the less hierarchical metabolic networks grow 2.25 times faster than those with more hierarchical networks (*p* < 0.0002).

**Figure 2.**
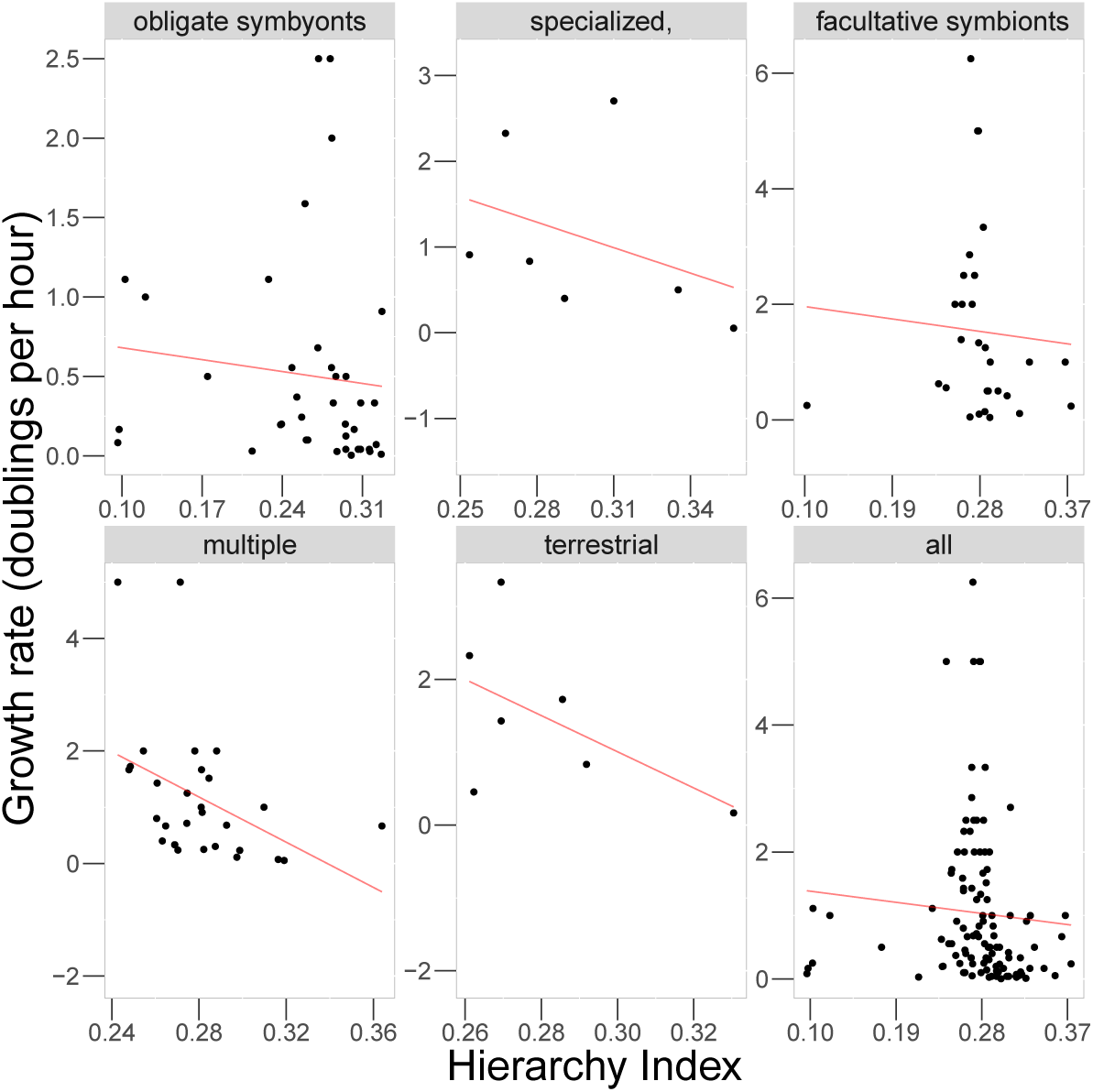
The relationship between hierarchical character and growth rate reflects fundamental tradeoffs between growth and yield and is informative about the ecological niche the bacteria occupy. Overall, growth rate is inversely correlated with hierarchy (Spearman’s rank correlation, *ρ* = −0.31, *p* = 0.00065). When controlling for bacterial environment the trend becomes stronger (*ρ* = −0.41, *p* = 0.0001). The outlier in the facultative parasite pane is *Borrelia burgdorferi*, which is an obligate parasite that alternates between insect and vertebrate hosts, and thus is similar to the obligate parasites. The particular strain also lacks a number of enzymes in its glycolysis pathway that are present in other *B. burgdorferi* strains that have hierarchy scores of 0.183 ± 0.002.

Thus the hierarchical character of the metabolic networks reflects the growth rate of the organisms and their environmental niche. These constraints of edge weight and tradeoffs between hierachical and ahierachical networks in metabolism are similar to those made in social networks and supply chains [20] [21].

### Relationship to other network properties

In addition to measuring hierarchy, we evaluated a number of other network statistics. We computed node count, edge count, modularity (as evaluated by the Girvan-Newman algorithm [29]), clustering coefficient, full diameter, effective diameter, number of strongly connected components, proportion of the nodes in the largest strongly connected component, and Luo Hierarchy score, an alternative metric of hierarchy that measures the proportion of edges that do not participate in any cycles. Edge and node count correlated most strongly with genetic distance. However, after these basic structural properties, the statistics that correlated most highly with genetic distance were the Girvan-Newman modularity score and the GRC hierarchy score (Table 1). We also computed the partial correlation for each variable with genetic distance, controlling for the others, and found that the GRC metric had the highest partial correlation.

**Table 1.**
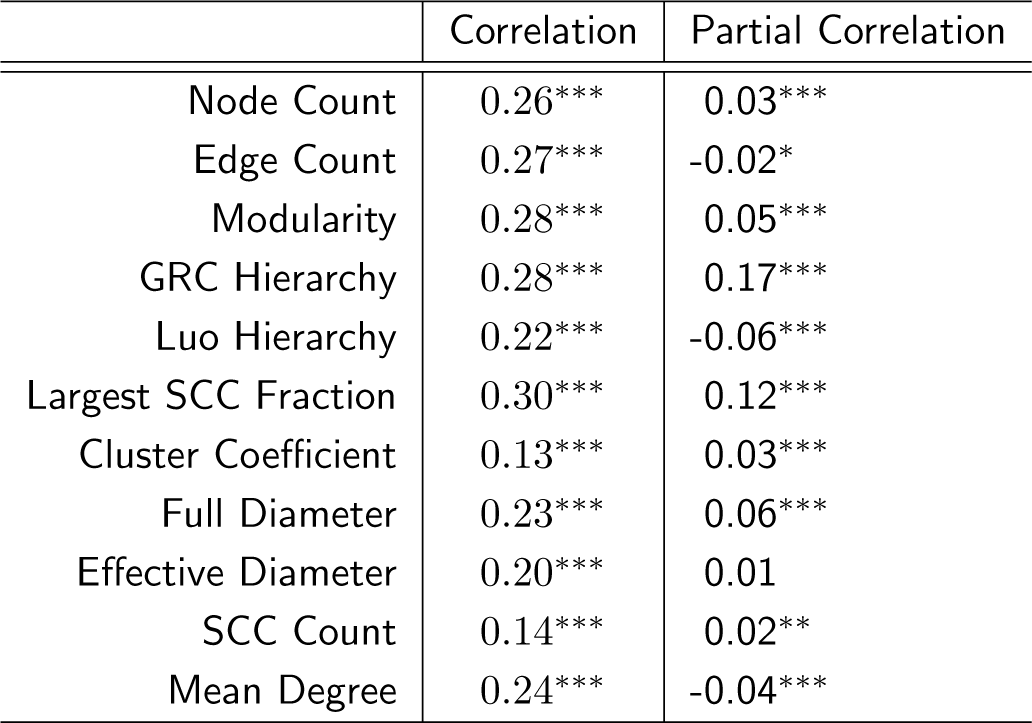
Correlation of network statistics with phylogenetic distances, and partial correlation of network statistic with phylogenetic distance, controlling for the other variables. The correlation and partial correlation of GRC Hierarchy metric with genetic distance is higher than all other non-trivial metrics. ^***^: *p* value < 0.001, ^**^: *p* value < 0.01, ^*^: *p* value < 0.05.

|  | Correlation | Partial Correlation |
| --- | --- | --- |
| Node Count | 0.26*** | 0.03*** |
| Edge Count | 0.27*** | -0.02* |
| Modularity | 0.28*** | 0.05*** |
| GRC Hierarchy | 0.28*** | 0.17*** |
| Luo Hierarchy | 0.22*** | -0.06*** |
| Largest SCC Fraction | 0.30*** | 0.12*** |
| Cluster Coefficient | 0.13*** | 0.03*** |
| Full Diameter | 0.23*** | 0.06*** |
| Effective Diameter | 0.20*** | 0.01 |
| SCC Count | 0.14*** | 0.02** |
| Mean Degree | 0.24*** | -0.04*** |

### Hierarchy Over Time

The hierarchy of the KEGG bacteria and reconstructed ancestors seems to first increase, and then decrease with distance from the root of the tree (Figure 3). Interestingly, with the dataset of 2,935 from the latest KEGG database, the correlation of modularity and distance from the root of the tree found by Kreimer *et al.* [17] is actually reversed. Modularity appears to increase rather than decrease with distance from the root, (Figure 4). This correlation remains positive when restricting analysis to the species used by Kreimer *et al.*.

**Figure 3.**
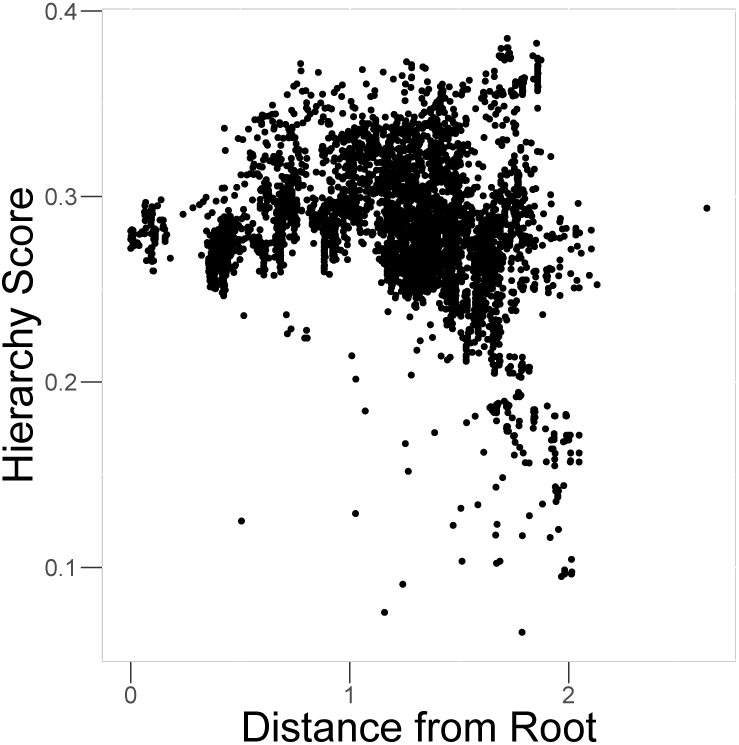
Hierarchy has a slight overall decrease with phylogenetic distance (Spearman’s rank correlation, *ρ* = −0.06, *p* < 10^−6^). Hierarchy appears to increase and then decrease further from the root of the tree.

**Figure 4.**
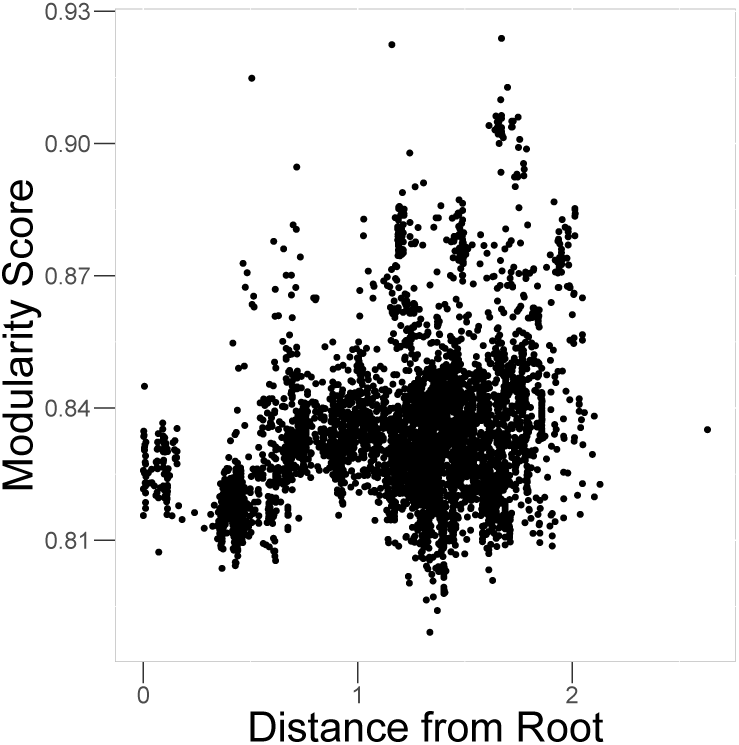
Modularity increases with phylogenetic distance (Spearman’s rank correlation, *ρ* = 0.31, *p* < 10^−15^).

As bacteria specialize to niches in a given ecosystem, they take on different metabolic strategies, which are reflected in the hierarchical profile of the metabolic network. This difference in strategies is consistent with the rise and fall of hierarchy over the evolutionary trajectory. As microbes first adapt to new environments or habitats (niche *sensu* Grinnell) they must gain novel metabolic functions, which are added as pathways in the periphery in the network and which increases the hierarchical character [30]. As complex relationships develop within the habitats, and bacteria adapt to different resource use profiles and competitive strategies (niche *sensu* Elton), the hierarchical profile of the metabolic network diversifies. Thus, the decrease in hierarchy over evolutionary time is caused by more bacteria specializing in a rapid-growth strategy, but the increasing variance in hierarchy reflects the fact that not all bacteria adopt this strategy. In studying the adaptive strategies chosen by different bacteria, we may be able to make inferences about the bacteria and their environments, as well as the interplay between evolutionary and ecological dynamics.

### Correlation of Modularity and Hierarchy

Hierarchy and modularity are global properties of metabolic networks. Both correlate with bacterial specialization and both change with distance from the root of the phylogenetic tree. Using the method of phylogenetic independent contrasts to look for correlation independent of phylogenetic structure, we found a moderate inverse correlation between modularity and hierarchy (Pearson correlation *r* = −0.18, *p* < 10^−15^), suggesting little evolutionary relationship between modularity and hierarchy [31]. Interestingly, simulated Boolean networks demonstrate a positive correlation between modularity and hierarchy [19].

## Conclusion

Characterizing the hierarchical structure of metabolic networks is useful in understanding the constraints under which these networks evolve. Hierarchy correlates with phylogenetic divergence, as would be expected for a trait subject to natural selection. This correlation is similar to the correlation of phylogenetic distance and modularity, suggesting that the hierarchical organization of networks, like modular organization, is important for function. However, modularity should be viewed as complementary to, rather than supplanted by, hierarchy when analyzing the global organization of metabolic networks. Both structural properties are conserved across phylogenies and evolve together. A better understanding of the character of metabolic networks is valuable in the growing field of ‘reverse ecology,’ in which the observed networks can be used to make inferences on possible environments [2][32][33].

By algorithmically reconstructing the metabolic networks, we are able to perform a larger-scale analysis than has previously been reported. Although the reaction annotations in KEGG may be prone to errors or omissions, we find that the GRC hierarchy metric is robust to small amounts of reaction omissions or reversals. By expanding the scope of the analysis, we find that modularity is actually inversely correlated with distance from the root of the tree, contrary to what has been found in previous studies of a more limited set of bacteria.

From reconstructed ancestral metabolic networks, we are able to infer how hierarchy evolves in networks over time, and understand the interplay between evolutionary and ecological dynamics. Hierarchy shows an increase followed by a decrease across the phylogenetic tree, which is reflective of the adaptive process of bacteria, first to novel fundamental niches, and then to a realized niche. The net trend in decreasing hierarchy reflects a dominance of fast-growth, low-efficiency strategy.

## Methods

### Hierarchy Metric

Hierarchy scores were calculated using the global reaching centrality metric developed by Mones *et al.*, which is based on the local reaching centrality [23]. The local reaching centrality (LRC) of a node in a network is the fraction of the nodes of the network that can be reached starting at the focal node. More precisely, if the metabolic network is represented as a graph, *G* = *(V, E)* it can be said that *v* reaches *v′* if there exists a series of edges (*v, v_i_*), (*v_i_, v_j_*)…(*v_j_, v′*) ∊ *E.* Let *R(v)* be the set of nodes *v′* ∊ *V* where *v′* is reachable from *v.* Then the LRC of *v* is 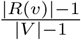. The GRC is then 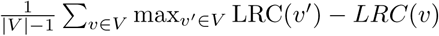

### Modularity Metric

The modularity metric was calculated using the SNAP package [34]. The modularity of a network is the optimal partitioning of the nodes into clusters to maximize 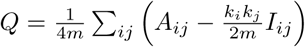. Where *m* is the number of edges in the network, *A_ij_* is the adjacency matrix, i.e. *A_ij_* is 1 if there is an enzyme that converts metabolite *i* into metabolite *j. k_i_* is the number of reactions that metabolite *i* participates in, and *I_ij_* = 1 if *i* and *j* are in the same module, and −1 otherwise. Since finding the global optimal of *Q* is an NP-hard problem, we use the method developed by Girvan and Newman, which partitions the network by iteratively removing the edge with the highest betweenness centrality [29].

### Robustness of Reconstruction

The KEGG database is large, with heavy manual curation; however, this does not mean that the data are always perfect. A reaction may be favorable in one direction in a model organism in laboratory conditions, but might proceed in the opposite direction or become bidirectional in different environments or species. It is also possible that reactions are missing from the database, or that an enzyme placed in an orthology group based on the study of one species may catalyze a different reaction in other species. To evaluate robustness to errors in the KEGG database, we examined the network for the well-studied bacterium, *E. coli.* We performed 100 replicates dropping or reversing 10% of the reactions, evaluated the hierarchy scores of these networks, and calculated the spread of the central 95% of hierarchy scores.

### Reconstruction of Genetic Distance

Following the methods often used in bacterial comparative genomics [35][3][36], for each of the 2,935 species, the 16s ribosomal sequence from KEGG was aligned to the Greenegenes database using PyNast, resulting in multiple sequence alignments for the 2,935 species [37][38]. The genetic distances between all pairs of bacterial species were computed using the Kimura distance metric [39].

### Reconstruction of Networks

For each bacterial species, a network of metabolites was inferred based on the enzymes present in the genome, the reactions known to be catalyzed by the enzymes present or orthologous enzymes, and a database of reaction substrates and products. The KEGG database of the genomic content of the 2,935 bacterial genomes was used to identify which enzyme classes were present in each genome and which reactions were present [40]. The reaction information from KEGG was supplemented by a the bioreaction database from Stelzer *et al.* which excludes currency metabolites, improves on predictions of directionality of reactions, and, for reactions with multiple substrates and products, provides carbon tracking of which substrates are converted to which products [41]. Using this reaction information, networks were constructed with metabolites as nodes, and a directed edge was placed between metabolites if there was a reaction that converted one metabolite to another. If reactions were reversible, then bi-directional edges were added between the substrates and products.

### Ancestral Networks

To construct the ancestral networks, a phylogenetic tree was reconstructed using RAxML 8.2.9 and the 16-state GTR nucleotide substitution model with gamma rate heterogeneity [42]. The branch-length weighted average bootstrap support of the partitions over 300 trees was 85.4. Using the maximum likelihood estimate of the best tree, at each interior node of the phylogenetic tree, a genome was constructed using the Fitch small parsimony algorithm. In cases where the presence or absence of a gene was equally parsimonious, the gene was randomly selected to be included. These ancestral genomes were then used to reconstruct ancestral networks, just as the networks were constructed on the leaves of the tree.

### Niche strategies

Growth rate data for 113 bacterial species and environmental annotations for those bacteria, which for 68 species were gathered from NCBI, and manual curation following literature review was used for the remaining 45 [43][4]. Due to their low number, the two aquatic species in the data set were excluded from further analysis. Correlations were calculated as Spearman’s *ρ.* To calculate correlation controlling for environment, *ρ* was calculated within each environment, and a species weighted-average across environments was computed. Due to several bacteria having the same growth rates, *p*-values were calculated using permutation tests rather than the Student’s t-distribution approximation. Significance tests were performed with 100,000 permutations each. For the overall *ρ*, permutations were done across all bacteria. To test the strength of the habitat-controlled correlation, growth rates were permuted within habitat classes and the species-weighted *ρ* was computed for each permutation. To test the effect of controlling for the environment, habitat labels were permuted and the difference between the species weighted *ρ* and overall *ρ* was computed.

## Consent to Publish

All authors have approved this manuscript for submission. This work has not been published or submitted elsewhere.

## Author Contributions

AG designed the experiment, carried out the analysis, and wrote the paper. MF provided critical guidence on the direction of the work, and revision of the manuscript.

## Availability of the Data

The dataset supporting the conclusions of this article, specifically the reconstructed metabolic network (Supplementary File 1) and topological statistics (Supplementary File 2) of these networks, are included within the article (and its additional files).

## Acknowledgments

We are grateful to B Callahan, E Borenstein and J Leskovec for helpful feedback with this work, and to editor Richard Goldstein and two anonymous reviewers for there insightful comments.

